# Dual NLRC4 and non-canonical inflammasome signaling drives human GSDMD-mediated killing of *Shigella flexneri* independently of bacterial cardiolipin

**DOI:** 10.64898/2026.01.11.698901

**Authors:** Mateusz Szczerba, Marcia B. Goldberg

## Abstract

Macrophages internalize and kill bacteria and thus are crucial for clearing bacterial infections. Although macrophage killing of some intracellular bacteria requires inflammasomes, the specific mechanisms of inflammasome-dependent killing are incompletely understood. Here we show that, upon infection with an intracellular pathogen *Shigella flexneri*, human macrophages activate a robust NLRC4-caspase-1 inflammasome response that restricts intracellular bacterial replication independently of pyroptosis. Gasdermin D (GSDMD) cleavage is required for bactericidal activity, revealing a GSDMD-dependent mechanism of bacterial killing independent of host cell death. We find that GSDMD-mediated killing of *S. flexneri* does not require bacterial cardiolipin, identifying a cardiolipin-independent mode of bacterial targeting. Priming macrophages with interferon (IFN)-γ enhances killing of intracellular *S. flexneri* by promoting involvement of caspase-4, which cooperates with caspase-1 to potentiate GSDMD function. These results identify a dual engagement of canonical and non-canonical inflammasomes that leads to macrophage killing *S. flexneri* while preserving host cell integrity. Furthermore, these findings uncover a previously unrecognized cardiolipin-independent mechanism of GSDMD-mediated bacterial killing with broad implications for immune cell control of cytosolic bacterial pathogens.

## INTRODUCTION

Macrophages are a crucial component of innate immunity against bacterial infections as they internalize and kill bacteria. Macrophage killing of bacteria occurs through diverse mechanisms, including phagolysosomal degradation, reactive oxygen and nitrogen species, and xenophagy.^1–3^ *Shigella* are cytosol-adapted pathogens that rapidly suppress many of these innate immune responses. *Shigella* escape the phagosome using type III secretion system effectors IpaB and IpaC, evading most phagolysosomal killing early during infection.^4–6^ Expression of autophagy antagonists, like IcsB, allow *Shigella* to escape xenophagy.^7^ *Shigella* also activate inflammasome signaling in infected macrophages,^8–10^ yet it is unclear to what degree inflammasomes contribute to macrophage control of *Shigella* or other bacteria.

Inflammasomes are cytosolic innate immune macromolecular complexes that assemble in response to infection and cellular stress.^11–16^ Inflammasome assembly is triggered by sensing of infection or danger signals by pattern recognition receptors, which activates caspase-1 or caspase-4. Activated caspases promote cleavage and secretion of pro-inflammatory cytokines interleukin-1β (IL-1β) and IL-18, as well as cleavage of the pore-forming protein gasdermin D (GSDMD), which leads to pyroptotic cell death.^17–21^ *Shigella* infection of human macrophages activates caspase-1 via the canonical NLRP3 and NLRC4 inflammasomes, and caspase-4 via the non-canonical NLRP11 inflammasome.^8–10^ Although inflammasome activation in human macrophages has been shown to be crucial for clearance of several intracellular bacterial pathogens, including *Salmonella*, *Legionella*, and *Burkholderia*, the specific mechanisms of inflammasome-mediated killing of bacteria are poorly understood,^22–25^ and whether inflammasomes contribute to killing of cytosolic *Shigella* in macrophages is unclear.

Here we investigated whether and how inflammasomes contribute to macrophage control of *Shigella*. We found that *Shigella flexneri* infection of human macrophages activates the NLRC4-caspase-1 inflammasome, leading to GSDMD-dependent bacterial killing that occurs independently of host pyroptosis. GSDMD executes a distinct bactericidal mechanism that restricts *S. flexneri* survival within intact macrophages. This GSDMD-mediated bactericidal activity is independent of cardiolipin, previously proposed as the essential lipid target for GSDMD pore formation on bacterial membranes.^17,19^ IFNγ priming of macrophages further enhances this response by upregulating caspase-4 and caspase-1, which amplifies GSDMD activation. Altogether, our study identifies dual engagement of canonical and non-canonical inflammasome signaling in macrophage anti-*Shigella* defenses that depend on a previously undescribed mechanism by which GSDMD directly targets intracellular pathogens.

## RESULTS

### Priming of macrophages with IFNγ restricts cytosolic *S. flexneri*

IFNγ is required to control intracellular *S. flexneri* in intestinal epithelial cells in mice in a manner that depends on natural killer (NK) cells.^26^ IFNγ, produced primarily by NK cells and T lymphocytes, is a cytokine that promotes activation of classical macrophages (M1), enhances macrophage antimicrobial effector functions,^27,28^ and primes inflammasomes.^29,30^ To begin to elucidate the role that macrophages play in host restriction of *S. flexneri*, we tested whether priming with IFNγ altered bacterial survival. Priming macrophages with IFNγ was associated with a significant decrease in accumulation of intracellular *S. flexneri*, as primed macrophages harbored nearly 10-fold fewer bacteria compared to unprimed macrophages (Figures 1A-1B and S1A), indicating that, as it does in mouse intestinal cells,^26^ IFNγ activates a human macrophage mechanism of *S. flexneri* restriction.

**Figure 1.**
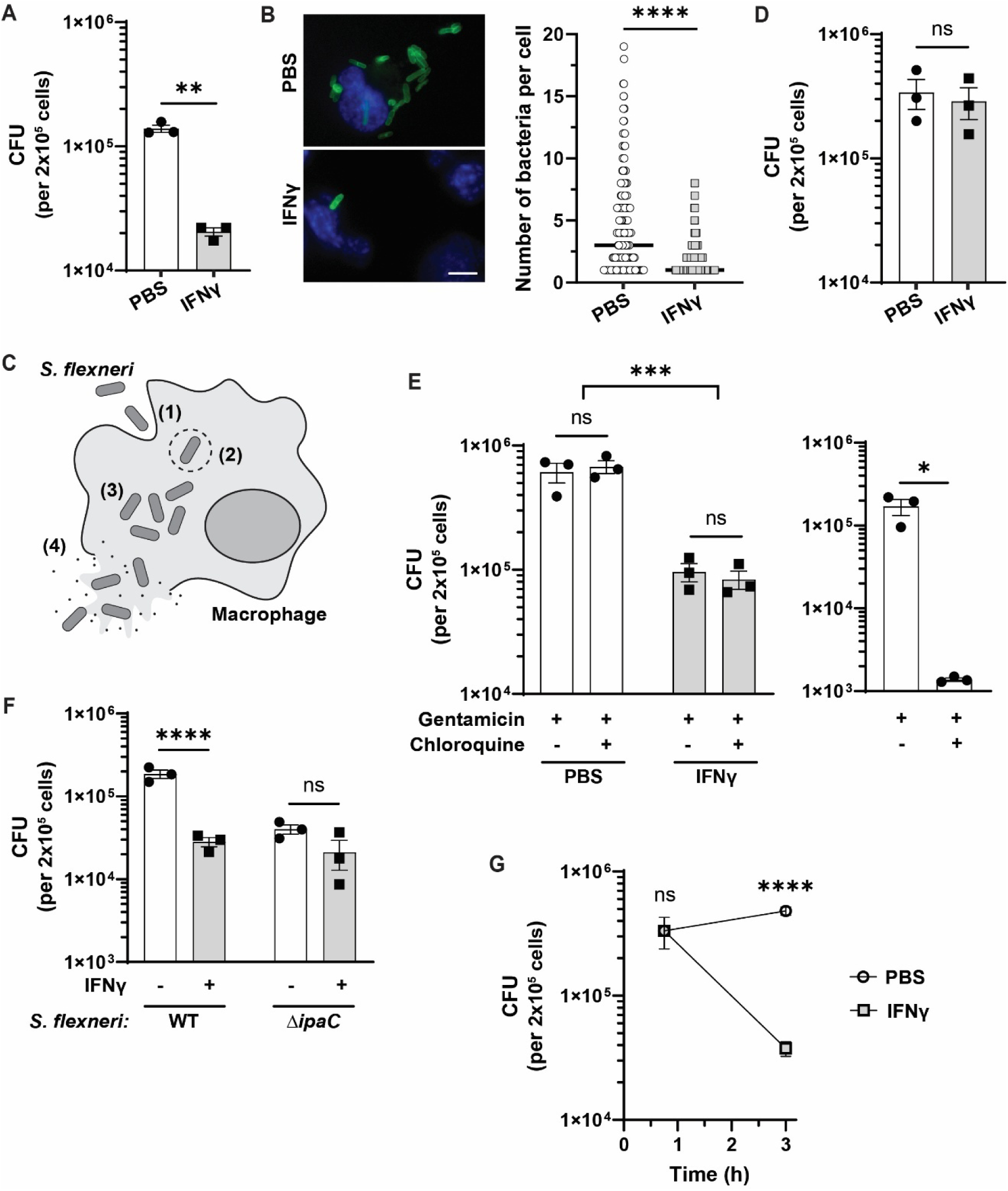
IFNγ priming of human macrophages leads to killing of *S. flexneri* in the cytosol. (A) Reduction of intracellular *S. flexneri* accumulation in IFNγ-primed macrophages at 3 hours of infection. (B) Reduction of bacterial numbers harbored by individual macrophages by IFNγ priming. Immunofluorescence at 2 hours of infection using Hoechst (blue) and antibody to *S. flexneri* (green) (representative images, left panel) and numbers of bacteria per cell (right graph). Each symbol represents one infected cell. Minimum of 360 cells (120 per biological replicate) were scored for each condition. Scale bar: 10 µm. (C) Schematic of *S. flexneri* infectious cycle in a human macrophage: (1) internalization by phagocytosis, (2) escape from the pathogen-containing vacuole, (3) replication in the cytosol, (4) release from the macrophage upon pyroptosis. (D) Absence of reduction of intracellular *S. flexneri* in IFNγ-primed macrophages at 45 minutes of infection. (E) Left graph, lack of impact of IFNγ priming on vacuolar escape. Right graph, *S. flexneri* Δ*ipaC*, unable to escape vacuoles, as a control for chloroquine activity. Gentamicin kills extracellular bacteria and chloroquine kills intravacuolar bacteria. (F) IFNγ restricts WT but not Δ*ipaC S. flexneri*. Bacterial counts, with and without IFNγ priming, in macrophages infected for 2 hours. (G) IFNγ activates a bactericidal mechanism against *S. flexneri*. Intracellular *S. flexneri* quantified in macrophages with and without IFNγ priming at the indicated times during infection. Data represent the mean ± SEM (A, D-G) or the median (B). \**p* < 0.05, \*\**p* < 0.01, \*\*\**p* < 0.001, \*\*\*\**p* < 0.0001, ns, not significant, by two-tailed unpaired Student’s t-test with Welch’s correction (A, D, right graph in E, G) or ordinary two-way ANOVA (left graph in E, F).

Upon internalization by macrophages, *S. flexneri* localizes to pathogen-containing vacuoles, which it rapidly escapes using the type III secretion system (Figure 1C), thereby releasing it into the cytosol, where it replicates.^4–6^ At 45 minutes after initial contact, at which point most infecting bacteria are internalized, we found no difference in numbers of viable intracellular bacteria between unprimed and primed macrophages (Figure 1D), indicating that the impact of IFNγ on *S. flexneri* survival and/or replication occurs post-internalization. To assess IFNγ impact on intravacuolar *S. flexneri*, we quantified bacterial numbers in the presence of chloroquine, which is bactericidal in acidic compartments such as the pathogen-containing vacuole.^31^ Numbers of *S. flexneri* were not reduced by chloroquine treatment of unprimed or IFNγ-primed macrophages (Figure 1E, left graph), indicating that IFNγ does not impair the ability of *S. flexneri* to escape the vacuole. In contrast, *S. flexneri* Δ*ipaC*, which is unable to escape the vacuole,^4^ was highly sensitive to chloroquine (Figure 1E, right graph). Furthermore, IFNγ-priming of macrophages had minimal impact on numbers of the Δ*ipaC* mutant (Figures 1F and S1B), indicating that IFNγ restriction occurs in the cytosol. The Δ*ipaC* mutant was recovered in lower numbers than wild-type *S. flexneri*, consistent with the Δ*ipaC* mutant residing in the vacuoles, where other mechanisms are known to clear bacteria.^32,33^ Altogether, these data demonstrate that IFNγ restriction occurs in the cytosol, after *S. flexneri* escapes from the vacuoles.

**IFN**γ priming results in killing *S. flexneri*

The effect of IFNγ priming of macrophages on intracellular *S. flexneri* could be bacteriostatic or bactericidal, each of which would result in reduced accumulation of bacteria. To discriminate between the two mechanisms, we assessed *S. flexneri* accumulation longitudinally during infection. We observed a significant decrease in bacterial numbers as the infection progressed in IFNγ-primed macrophages (Figure 1G), which is consistent with IFNγ priming activating a bactericidal mechanism that kills intracellular *S. flexneri*.

### The cytosolic killing of *S. flexneri* is independent of pyroptosis and macrophage lysis

As inflammasome activation leads to activation of pyroptosis, we tested whether macrophage cell death *per se* is responsible for the observed IFNγ-dependent killing of *S. flexneri*, as has been described for other pathogens.^22–25^ We found that IFNγ priming does not impact *S. flexneri*-induced macrophage death (Figure 2A). To directly test whether macrophage lysis contributes to *S. flexneri* restriction, we inhibited plasma membrane rupture using glycine (Figure 2B).^34^ Although treatment of macrophages with glycine led to inhibition of macrophage death (Figure S2A), glycine had no impact on restriction of intracellular *S. flexneri* (Figure 2C), indicating that restriction does not depend on macrophage lysis. Similarly, treatment of macrophages with disulfiram, an inhibitor of gasdermin D (GSDMD) membrane pore formation,^35^ led to inhibition of pyroptosis as measured by macrophage death (Figure S2B) and release of cleaved caspases (Figure 2D), yet had no impact on numbers of intracellular *S. flexneri* (Figure 2E). These data indicate that macrophage killing of *S. flexneri* is independent of pyroptosis.

**Figure 2.**
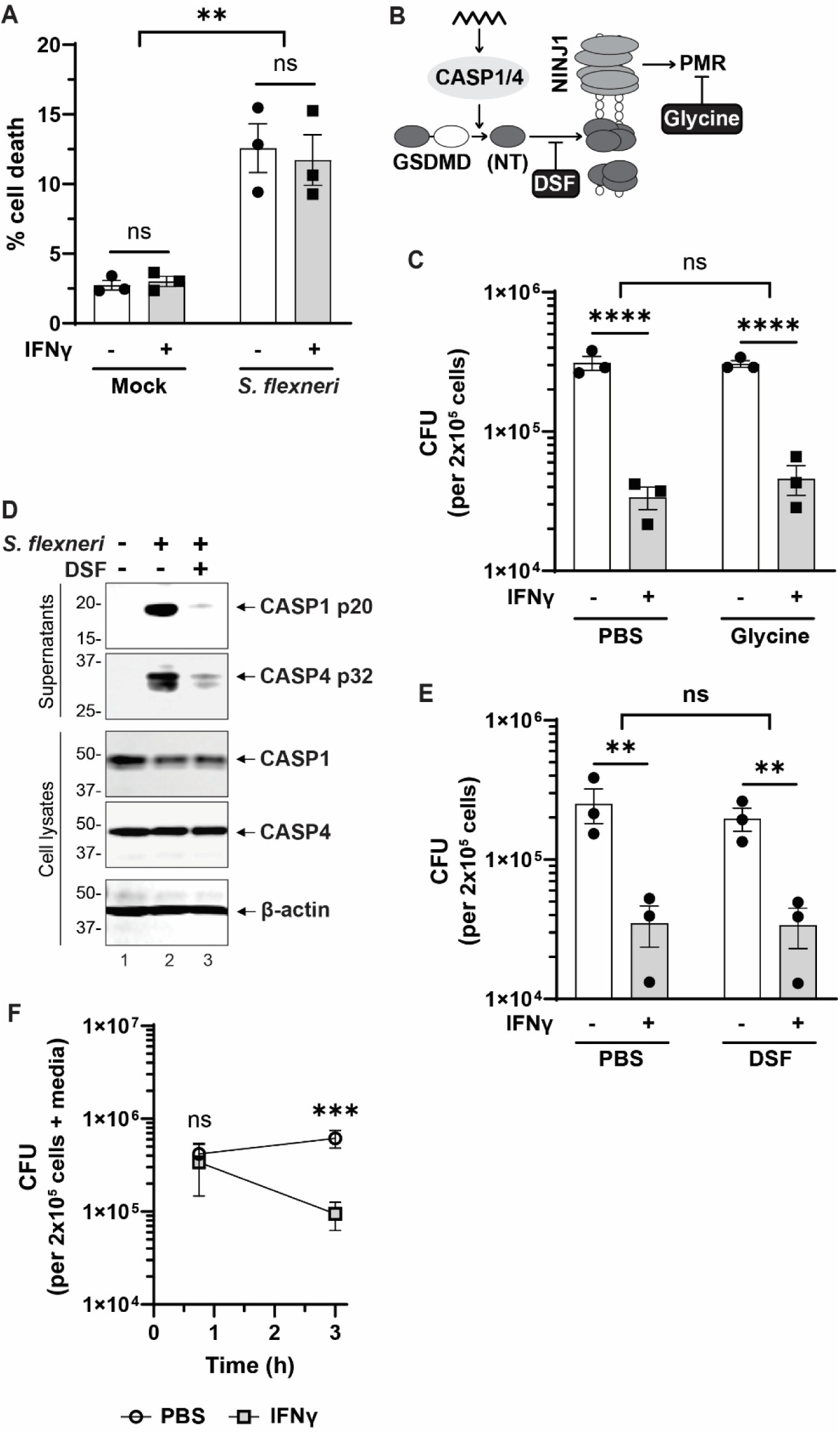
IFNγ-mediated killing of *S. flexneri* occurs via an intracellular mechanism independent of host cell lysis. (A) Lack of impact of IFNγ priming on cell death of *S. flexneri*-infected macrophages. Cell death measured as lactate dehydrogenase release. (B) Schematic of pyroptosis with indicated sites of action of inhibitors of gasdermin D (GSDMD) N-terminal domain (NT) pore formation (DSF, disulfiram) and of ninjurin-1 (NINJ1) oligomerization and plasma membrane rupture (PMR; glycine). CASP1/4, caspase-1 and/or -4. (C) Intracellular *S. flexneri* upon inhibition of lytic cell death. Macrophages, primed or not primed with IFNγ, were treated with glycine in the absence of gentamicin. (D) Inhibition of *S. flexneri* infection-induced release of activated caspase-4 (CASP4 p32) and activated caspase-1 (CASP1 p20) by disulfiram. Representative immunoblots. (E) Intracellular *S. flexneri* upon inhibition of GSDMD plasma membrane pore formation. Macrophages, primed or not primed with IFNγ, were treated with disulfiram in the absence of gentamicin. (F) IFNγ mediated restriction is not due to bacterial loss in cell culture supernatant or detached macrophages. *S. flexneri* in cell culture supernatants and washes (which contain detached macrophages and released bacteria), and in attached macrophages. Samples collected at indicated times during infection. Graphed are bacterial counts combined for supernatants, washes, and attached cells. Data represent the mean ± SEM. \*\**p* < 0.01, \*\*\**p* < 0.001, \*\*\*\**p* < 0.0001, ns, not significant, by two-tailed unpaired Student’s t-test (F) or ordinary two-way ANOVA (A, C, E).

Although priming macrophages with IFNγ does not lead to increased pyroptosis, it could perturb cellular homeostasis in a manner that results in increased macrophage detachment from the tissue culture plates used for these assays, which might artifactually lower bacterial counts. To test this, we performed infections without gentamicin in the media and quantified *S. flexneri* in cell culture supernatants and washes (Figure 2F, “media”); this media fraction contains macrophages that have detached from the plate as well as bacteria released from cells. Even when including bacteria recovered from this fraction in the quantification, IFNγ priming was associated with a decrease in *S. flexneri* numbers (Figure 2F). Indeed, fewer bacteria were recovered from cell culture supernatants and washes of IFNγ-primed macrophages than of unprimed macrophages (Figure S3A). Altogether, these data demonstrate that killing of *S. flexneri* occurs via a cytosolic mechanism that is independent of host cell lysis.

### Intracellular restriction depends on caspase-1-mediated activation of GSDMD

To directly investigate whether inflammasomes contribute to restricting *S. flexneri*, we quantified intracellular bacteria in macrophages deficient in *GSDMD* (Figure S3A), which encodes the common terminal effector of inflammasome-mediated pyroptosis.^18,20^ Knocking out *GSDMD* was associated with a greater than 3-fold rescue of *S. flexneri* accumulation compared to wild-type (WT) macrophages (Figure 3A), indicating that GSDMD is a critical factor in restricting *S. flexneri*. Bacterial accumulation was rescued to a similar extent in both unprimed and IFNγ-primed macrophages (Figure 3A). This rescue is not due to release of fewer bacteria by *GSDMD*^-/-^ macrophages, as similar numbers or more bacteria were present in cell culture supernatants of *GSDMD*^-/-^macrophages than WT macrophages (Figure S3B). As for WT macrophages, IFNγ-primed *GSDMD*^-/-^ macrophages harbor fewer bacteria than unprimed *GSDMD*^-/-^ macrophages (Figure 3A), indicating the presence of an IFNγ-dependent GSDMD-independent mechanism of bacterial restriction; such mechanisms have been described previously^36,37^ and are not the focus of this work.

**Figure 3.**
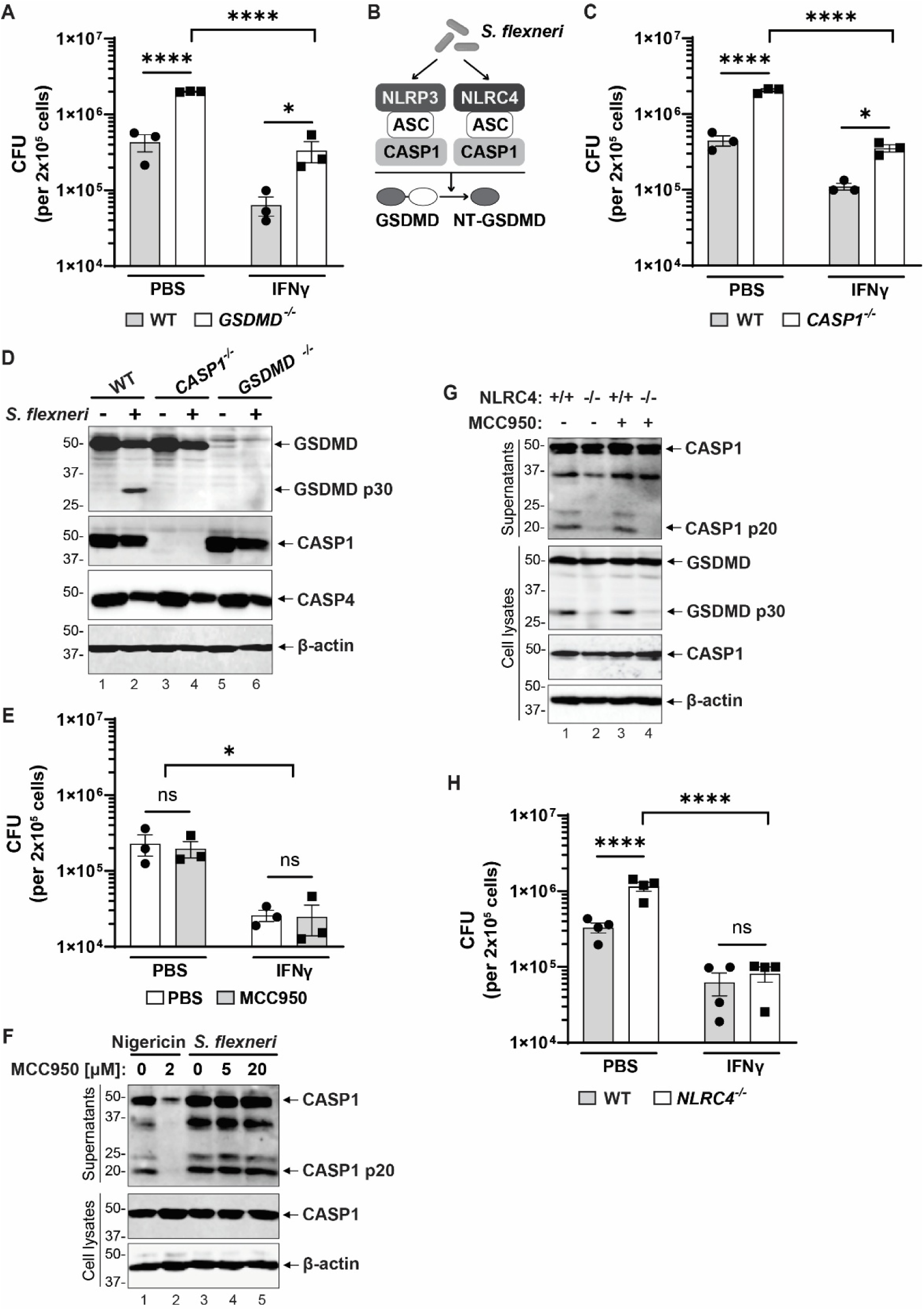
Restriction of *S. flexneri* in unprimed macrophages is mediated by the NLRC4/caspase-1/GSDMD inflammasome. (A) Rescue of intracellular *S. flexneri* in the absence of *GSDMD*, with or without IFNγ priming. (B) Schematic of *S. flexneri* activation of caspase-1 inflammasomes in human macrophages. ASC, apoptosis-associated speck-like protein containing a CARD; CASP1, caspase-1; NT-GSDMD, N-terminal pore-forming domain of GSDMD. (C) Rescue of intracellular *S. flexneri* in the absence of *CASP1*, with or without IFNγ priming. (D) Caspase-1 is required for *S. flexneri* infection-induced activation of GSDMD (GSDMD p30). Unprimed macrophages. (E-F) Lack of impact of NLRP3 inhibition by MCC950 on numbers of intracellular *S. flexneri* in macrophages, with or without IFNγ priming (E) and on infection-induced caspase-1 activation (F). Positive control: treatment with NLRP3 agonist nigericin. (G) Reduced processing of caspase-1 (CASP1 p20) and GSDMD (GSDMD p30) during *S. flexneri* infection in the absence of *NLRC4*. (H) Rescue in *NLRC4*^-/-^ macrophages of intracellular *S. flexneri* in the absence but not the presence of IFNγ priming. Immunoblots are representative (D, F-G). Data represent the mean ± SEM. \**p* < 0.05, \*\*\*\**p* < 0.0001, ns, not significant, by ordinary two-way ANOVA.

During infection of macrophages, *S. flexneri* activates GSDMD via two caspase-1 pathways (Figure 3B) and a caspase-4 pathway (Figure 4A).^8–10^ Infection of *CASP1*^-/-^ macrophages was associated with a greater than 3-fold rescue of bacterial counts (Figures 3C and S3A), both in unprimed and IFNγ-primed macrophages. This level of rescue of bacterial counts was similar to that observed in *GSDMD*^-/-^ macrophages (Figure S3C), suggesting that caspase-1 activation of GSDMD is the primary pathway restricting *S. flexneri*. This is consistent with the observation that knocking out *CASP1* inhibits *S. flexneri*-induced activation of GSDMD (Figure 3D). These data indicate that caspase-1-mediated activation of GSDMD contributes significantly to the restriction of intracellular *S. flexneri*.

**Figure 4.**
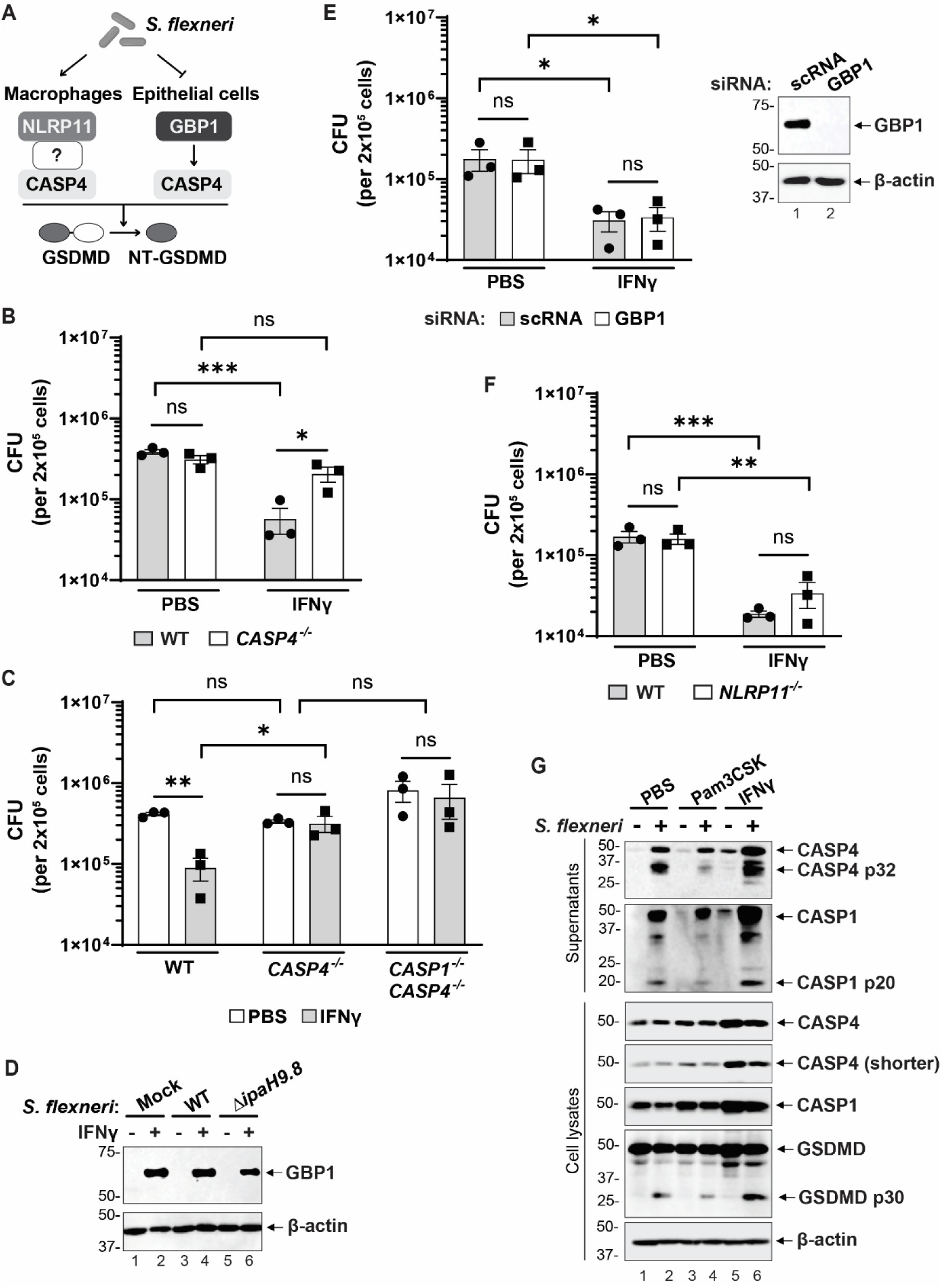
IFNγ-dependent increases in caspase-4 and caspase-1 are associated with enhanced bacterial restriction independent of GBP1 and pattern-recognition receptors. (A) Schematic of *S. flexneri* activation of inflammasomes that depend on caspase-4 (CASP4). In macrophages, *S. flexneri* infection activates GSDMD via NLRP11.^9^ In epithelial cells, caspase-4 is activated by guanylate-binding protein 1 (GBP1) recognition of LPS, which *S. flexneri* inhibits.^43–45^ NT-GSDMD, N-terminal pore-forming domain of GSDMD. (B-C) The absence of *CASP4* (B) or *CASP4* and *CASP1* (C) impacts intracellular *S. flexneri* in the presence but not absence of IFNγ-priming. (D) Levels of GBP1 in macrophages infected with WT or Δ*ipaH9.8 S. flexneri*. (E) Lack of impact of GBP1 on restriction of *S. flexneri*. GBP1 depletion by RNA interference (siRNA). scRNA, scrambled RNA interference control. (F) Lack of significant impact of NLRP11 on intracellular *S. flexneri* in macrophages, with or without IFNγ priming. (G) IFNγ priming of *S. flexneri* infection is associated with increased levels of cellular caspase-4 (CASP4) and caspase-1 (CASP1), increased release of cleaved CASP4 (CASP4 p32) and cleaved CASP1 (CASP1 p20) into the cell culture supernatants, and increased cellular cleavage of GSDMD to release the pore-forming domain (GSDMD p30). Shorter, shorter exposure. Immunoblots are representative (D-E, G). Data represent the mean ± SEM. \**p* < 0.05, \*\**p* < 0.01, \*\*\**p* < 0.001, ns, not significant by ordinary two-way ANOVA.

### Caspase-1-mediated restriction of intracellular *S. flexneri* in unprimed macrophages depends on NLRC4

During *S. flexneri* infection of macrophages, caspase-1 activation occurs downstream of both NLRP3 and NLRC4 (Figure 3B).^8,10^ Inhibition of NLRP3-mediated activation of caspase-1 with the inhibitor MCC950^38^ did not rescue intracellular *S. flexneri* (Figure 3E), indicating that NLRP3 is not required for restriction. Consistent with the lack of rescue with MCC950, caspase-1 remained activated in its presence, even at concentrations ten-fold higher than what is required for inhibition of NLRP3 activation by the NLRP3 agonist nigericin^39,40^ (Figure 3F). These data indicate that NLRP3 is not required for caspase-1 restriction of *S. flexneri*.

In contrast, in unprimed macrophages, deletion of *NLRC4* resulted in markedly reduced activation of caspase-1 and GSDMD (Figure 3G) and in a greater than 3-fold increase in accumulation of intracellular *S. flexneri* (Figure 3H), demonstrating that the NLRC4/caspase-1/GSDMD pathway is required for killing of intracellular *S. flexneri*. NLRP4 is known to recognize components of the type III secretion system,^41,42^ which is present in *S. flexneri* and many other gram-negative bacterial pathogens. Upon IFNγ priming, *NLRC4*^-/-^ macrophages harbored bacterial numbers similar to those in WT macrophages (Figure 3H), indicating that NLRC4 contributes to *S. flexneri* killing only in the absence of IFNγ priming.

### Efficient IFNγ-dependent killing of *S. flexneri* depends on caspase-4

In addition to activating the NLRP3 and NLRC4 inflammasomes, *S. flexneri* infection activates the non-canonical inflammasome mediated by caspase-4 (Figure 4A).^9^ Using *CASP4*^-/-^ macrophages (Figure S3B), we found that in the absence of priming, caspase-4 was not required for restriction of *S. flexneri*, but in IFNγ-primed macrophages, knocking out *CASP4* was associated with increased accumulation of intracellular *S. flexneri* to numbers close to those recovered from WT macrophages (Figure 4B), indicating that caspase-4 is required for IFNγ-mediated killing of *S. flexneri*. Under these same IFNγ priming conditions, deletion of *CASP1* in *CASP4*^-/-^ macrophages was associated with enhanced rescue of *S. flexneri* (Figure 4C), demonstrating an additive effect of the two caspases. These data indicate that, upon IFNγ priming, both caspases contribute to *S. flexneri* restriction. They further suggest that activation of GSDMD downstream of caspase-4 occurs, at least in part, via activation of caspase-1. As for WT macrophages and macrophages described above that contain other specific deletions of inflammasome components, the efficiency of *S. flexneri* invasion into *CASP4*^-/-^ macrophages was unchanged by the deletion or IFNγ priming (Figure S3D).

### Caspase-4-mediated bacterial restriction is independent of GBP1

In human epithelial cells, guanylate-binding proteins (GBPs) activate caspase-4 by enhancing extraction of lipopolysaccharide (LPS) from gram-negative bacteria, facilitating LPS recognition and binding by caspase-4.^43,44^ Also in epithelial cells, *S. flexneri* type III secretion effector IpaH9.8 degrades GBP1,^45,46^ the most upstream protein of the GBP cascade. We found that in human macrophages, deletion of *S. flexneri* IpaH9.8 did not result in increased levels of GBP1 (Figure 4D), demonstrating that, in contradistinction to epithelial cells, IpaH9.8 does not degrade GBP1 in these immune cells. Furthermore, *S. flexneri* Δ*ipaH9.8* was restricted intracellularly similarly to WT *S. flexneri* (Figure S4A). Knock-down of GBP1 resulted in no change in intracellular *S. flexneri* (Figure 4E), demonstrating that, in human THP-1 macrophages, GBP1 is not required for control of intracellular bacteria, including for IFNγ-dependent caspase-4-mediated bacterial restriction.

### IFNγ priming increases levels of caspases and activation of GSDMD

NLRP11 is a pattern-recognition receptor required for efficient activation of caspase-4-mediated pyroptosis during *S. flexneri* infection of unprimed human macrophages (Figure 4A).^9^ Nevertheless, deletion of *NLRP11* had minimal impact on accumulation of intracellular *S. flexneri* (Figure 4F and S4B), demonstrating that IFNγ-activated caspase-4-mediated control of *S. flexneri* is primarily independent of NLRP11. Similarly, in IFNγ-primed macrophages, caspase-4- and caspase-1-mediated restriction of intracellular *S. flexneri* was independent of any of the tested pattern-recognition receptors and GBP1 (Figures 3 and 4).

IFNγ priming was associated with increased levels of caspase-4 and caspase-1, as well as increased activation of GSDMD compared to unprimed macrophages or macrophages treated with PAM3CSK, a TLR2/TLR1 ligand^47^ (cell lysates in Figure 4G, quantified in Figure S4C). IFNγ priming was also associated with increased release of activated caspase-1 and caspase-4 from *S. flexneri*-infected macrophages (supernatants in Figure 4G, quantified in Figure S4C). These results are most consistent with IFNγ-dependent increases in caspase-4 and caspase-1 being responsible for both the enhanced GSDMD activation and the increased intracellular bacterial control that occur during *S. flexneri* infection upon priming with IFNγ.

### Macrophage killing of *S. flexneri* is mediated by GSDMD

As restriction of intracellular *S. flexneri* required GSDMD (Figure 3) and enhanced killing of *S. flexneri* in IFNγ-primed macrophages was associated with enhanced activation of GSDMD (Figure 4), we examined how GSDMD contributes to bacterial killing. In *GSDMD*^-/-^ macrophages, replication of *S. flexneri* over time was significantly increased compared to WT macrophages (Figure 5A), and the number of bacteria per cell was greater for *GSDMD*^-/-^ macrophages than for WT macrophages (Figures 5B and S5A). Consistent with our finding that bacterial killing occurs in the cytosol (Figure 1), knockout of *GSDMD* had minimal impact on survival of *S. flexneri* Δ*ipaC* (Figure 5C), which remains in the pathogen-containing vacuole^4^.

**Figure 5.**
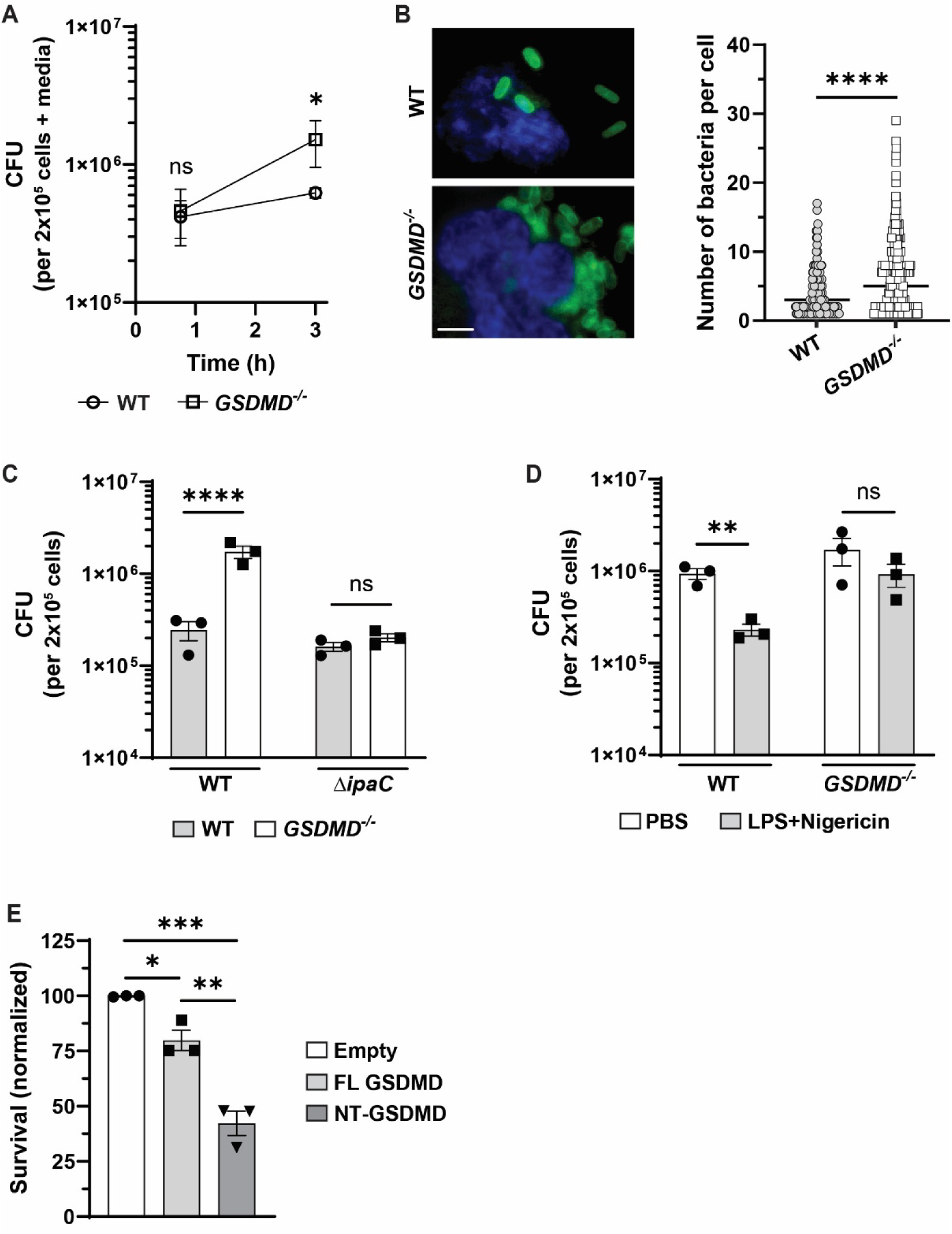
Inflammasome-dependent killing of *S. flexneri* is mediated by GSDMD. (A) Replication of *S. flexneri* in *GSDMD*^-/-^ macrophages. Numbers of *S. flexneri* in cell culture supernatants and washes (containing detached macrophages and released bacteria) and attached macrophages, at the indicated infection times. Graphed are bacterial counts combined for supernatants, washes, and attached cells. (B) Infected *GSDMD*^-/-^ macrophages harbor significantly more *S. flexneri* per cell than infected WT macrophages. Left panel, immunofluorescence at 2 hours of infection using Hoechst (blue) and antibody to *S. flexneri* (green) (representative images). Right graph, numbers of bacteria per cell. Each symbol in the scatter blot represents one infected cell. Minimum of 480 cells (160 per biological replicate) were scored for each condition. Scale bar: 10 µm. (C) Lack of impact of GSDMD on survival of intravacuolar *S. flexneri* Δ*ipaC*. (D) Enhanced GSDMD-dependent killing of intracellular *S. flexneri* in macrophages treated with extracellular LPS and nigericin. (E) Killing of intracellular *S. flexneri* upon infection of HEK293T cells expressing full-length GSDMD (FL GSDMD) or the GSDMD N-terminal pore-forming domain (NT-GSDMD). Survival is calculated using bacterial counts combined for supernatants, washes, and attached cells. Data represent the mean ± SEM (A, C-E) or the median (B). \**p* < 0.05, \*\**p* < 0.01, \*\*\**p* < 0.001, \*\*\*\**p* < 0.0001, ns, not significant, by two-tailed unpaired Student’s t-test (A-B) or ordinary one- (E) or two-way (C-D) ANOVA.

Treatment of macrophages with nigericin and extracellular LPS, which increases the activation of GSDMD,^19^ was associated with a reduction in intracellular *S. flexneri* (Figure 5D); this was not due to increased bacterial loss into the media from nigericin-induced cell death (Figure S5B). In macrophages lacking GSDMD, the nigericin-induced enhancement of bacterial killing was reduced (Figure 5D), indicating that this phenotype is largely GSDMD-dependent.

To test whether GSDMD-dependent killing of *S. flexneri* requires other inflammasome components, we expressed GSDMD in HEK293T cells (Figure S5C), which lack all inflammasome proteins. Expression of either full length GSDMD or the isolated N-terminal pore-forming domain (NT-GSDMD) was associated with significant decreases in intracellular *S. flexneri* (Figure 5E), indicating that GSDMD kills *S. flexneri* independently of other inflammasome components. NT-GSDMD was more efficient in killing of *S. flexneri* than the full-length protein (Figure 5E). Thus, NT-GSDMD is the major mediator of inflammasome-dependent killing of *S. flexneri*. Expression of GSDMD constructs had no impact on bacterial invasion of HEK293T cells (Figure S5D).

### GSDMD kills intracellular *S. flexneri* independently of oxidative stress

Because GSDMD forms pores in mitochondria,^48^ leading to release of mitochondrial reactive oxygen species that have bactericidal effects,^3^ we tested whether oxidative stress was required for GSDMD-mediated killing of *S. flexneri*.

Treatment with glutathione, an oxidative stress inhibitor,^49,50^ had minimal impact on GSDMD-dependent killing of intracellular *S. flexneri* (Figure S6A). This is consistent with our findings showing that treatment with disulfiram, which inhibits GSDMD-driven mitochondrial damage,^48^ did not impact macrophage restriction of *S. flexneri* (Figure 2E). These results indicate that mitochondrial damage and oxidative stress do not significantly contribute to GSDMD-mediated killing of *S. flexneri*.

### GSDMD-mediated killing of *S. flexneri* does not require bacterial cardiolipin

*In vitro*, GSDMD binds negatively charged lipids, including cardiolipin, a phospholipid found in bacterial and mitochondrial membranes.^17,19,51^ Among the three major lipids in enterobacterial membranes – phosphatidyl-ethanolamine, phosphatidyl-glycerol, and cardiolipin – cardiolipin is the only one to which GSDMD binds *in vitro*. Consequently, cardiolipin has been hypothesized to be the GSDMD ligand on bacteria required for direct bacterial killing.^17,19^

*S. flexneri* Δ*clsA*, which lacks cardiolipin as a result of deletion of the cardiolipin synthase A (ClsA),^52^ was restricted in human macrophages in a manner that was no different from WT *S. flexneri*. Deletion of *GSDMD* led to rescue of the intracellular Δ*clsA* mutant to the same levels as WT *S. flexneri*, and killing of the Δ*clsA* mutant was enhanced by IFNγ to the same degree as that of WT *S. flexneri* (Figure 6A). As for WT *S. flexneri* (Figure 5B), infected *GSDMD*^-/-^macrophages harbored more *S. flexneri* Δ*clsA* than WT macrophages (Figures 6B and S6B). And as for WT *S. flexneri* (Figure 5D), chemical activation of NLRP3 by treatment with nigericin and extracellular LPS was associated with a reduction of intracellular *S. flexneri* Δ*clsA* (Figure 6C). This nigericin-induced killing was independent of bacterial release into the media (Figure S6C), but dependent on GSDMD (Figure S6D); the extent of increased killing was similar for WT and Δ*clsA S. flexneri* (Figure S6E). Expression of NT-GSDMD in HEK293T cells infected with *S. flexneri* Δ*clsA* was associated with significant reduction of intracellular bacterial counts (Figure 6D), as observed for WT *S. flexneri* (Figures 5E and 6E), with no impact on *S. flexneri* Δ*clsA* invasion of HEK293T cells (Figure S5D). Moreover, in macrophages, as early as 50 minutes of infection, GSDMD co-localized with both WT *S. flexneri* and the Δ*clsA* mutant (Figure 6F) and did so in a manner that was not impacted by priming with IFNγ (Figure S6F). The Δ*clsA* mutant grows normally *in vitro* ^52^ and invaded macrophages as efficiently as WT *S. flexneri* (Figure S6G). Although reported to form filaments in the cytosol of epithelial cells at later stages of infection,^52^ at the times tested, the Δ*clsA* mutant maintained a bacillus morphology, resembling that of WT *S. flexneri* (Figure 6B, compared to 1B and 5B).

**Figure 6.**
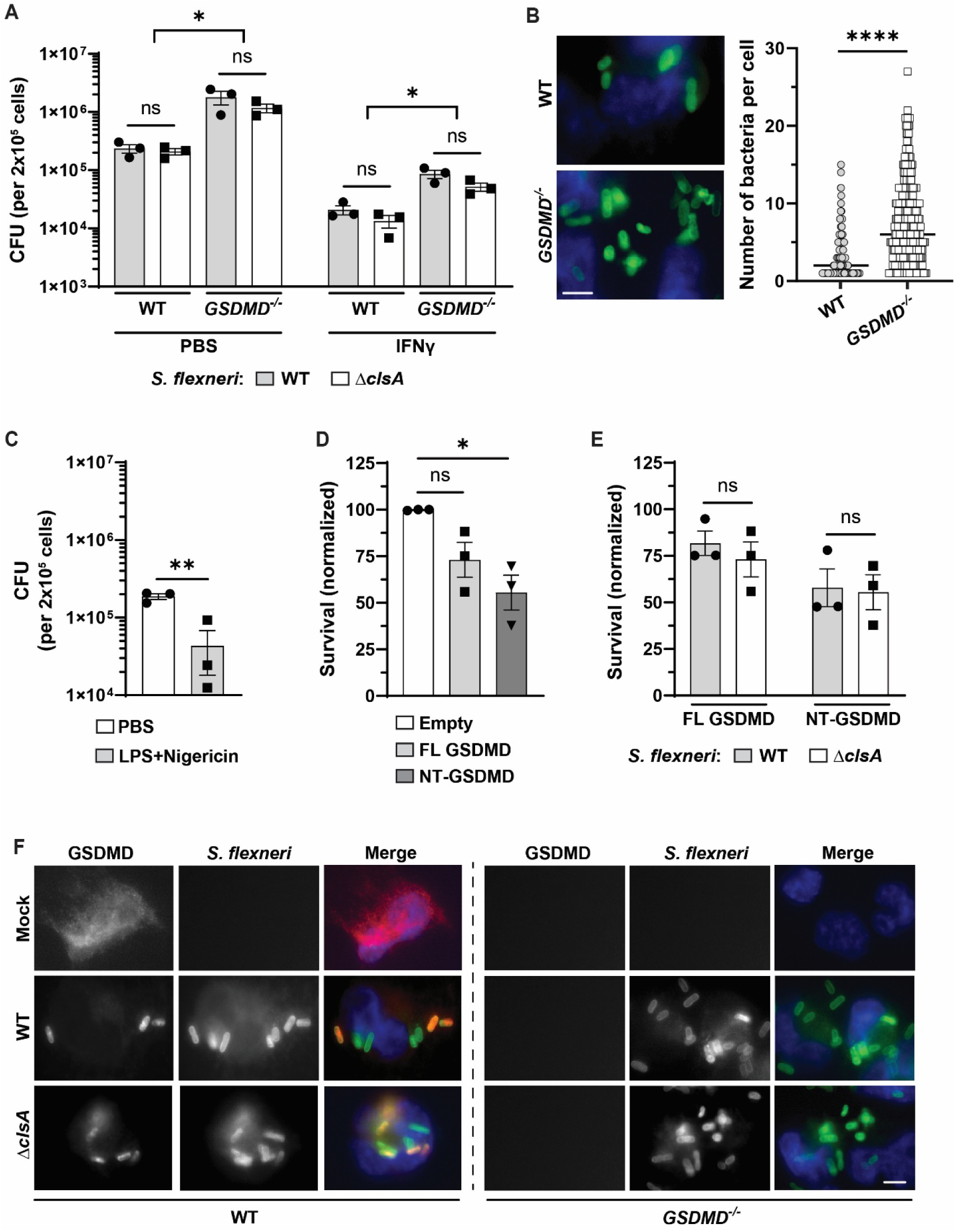
GSDMD kills intracellular *S. flexneri* independently of bacterial cardiolipin. (A) Lack of impact of cardiolipin on GSDMD-mediated killing, with or without IFNγ priming. WT and *GSDMD*^-/-^ macrophages infected with WT or cardiolipin mutant (Δ*clsA*) of *S. flexneri*. (B) *S. flexneri* Δ*clsA*-infected *GSDMD*^-/-^ macrophages harbor significantly more bacteria per cell than *S. flexneri* Δ*clsA*-infected WT macrophages. Left panel, immunofluorescence at 2 hours of infection using Hoechst (blue) and antibody to *S. flexneri* (green) (representative images). Right graph, numbers of bacteria per cell. Each symbol in the scatter blot represents one infected cell. Minimum of 420 cells (140 per biological replicate) were scored for each condition. Scale bar: 10 µm. (C) Enhanced killing of intracellular *S. flexneri* Δ*clsA* in macrophages treated with extracellular LPS and NLRP3 agonist nigericin. (D) Killing of intracellular *S. flexneri* Δ*clsA* upon infection of HEK293T cells expressing full-length GSDMD (FL GSDMD) or GSDMD N-terminal pore-forming domain (NT-GSDMD). Survival is calculated using bacterial counts combined for supernatants, washes, and attached cells. (E) Lack of significant differences in survival between WT and Δ*clsA S. flexneri* upon infection of HEK293T cells expressing indicated GSDMD constructs. For each bacterial strain, survival is calculated as compared to HEK293T cells transfected with the empty vector. (F) Colocalization of GSDMD and *S. flexneri*. Immunofluorescence using antibodies to GSDMD (red) and *S. flexneri* (green). Unprimed WT and *GSDMD*^-/-^ macrophages infected with WT or Δ*clsA S. flexneri* for 50 mins. Representative images. Data represent the mean ± SEM (A, C-D) or median (B). \**p* < 0.05, \*\**p* < 0.01, **** *p* < 0.0001, ns, not significant, by two-tailed unpaired Student’s t-test (B-C) or ordinary one- (D) or two-way (A and E) ANOVA.

Altogether, these data demonstrate that GSDMD kills intracellular *S. flexneri* via a mechanism that is independent of bacterial cardiolipin, indicating that GSDMD targeting of *S. flexneri* does not depend on cardiolipin or on its direct binding to any of the major bacterial membrane lipids.

## DISCUSSION

We have found for the first time that GSDMD mediates killing of a bacterial pathogen in the cytosol of macrophages. We found that GSDMD co-localizes with *S. flexneri* during infection of human macrophages (Figures 6F and S6F) and that macrophages kill *S. flexneri* in a manner dependent on GSDMD (Figures 3A and 5A-5D). Inflammasomes have been implicated in antimicrobial defense against certain bacterial pathogens,^22–25^ yet the mechanisms for restricting bacterial survival described to date rely on secondary mediators following pyroptosis, such as neutrophil extracellular traps and pore-induced intracellular traps.^23,54^ Whereas GSDMD is most extensively described for executing pyroptotic cell death by insertion of pores in the plasma membrane,^18,20^ we found that GSDMD kills *S. flexneri* in the cytosol of human macrophages (Figure 5C) independently of macrophage cell death (Figures 2 and S3B). Hence, we definitively identify a cytosolic effector of macrophage-intrinsic bacterial killing and show that inflammasomes act cell-autonomously to eliminate cytosolic bacteria.

An unexpected finding is that GSDMD killing does not require bacterial cardiolipin. As cardiolipin is the only major bacterial lipid shown to bind GSDMD, it was previously proposed as the essential lipid target for GSDMD pore formation on bacterial membranes.^17,19^ In human macrophages, we found that GSDMD co-localized with cardiolipin-deficient (Δ*clsA*) *S. flexneri*, as it did with WT *S. flexneri*, (Figures 6F and S6F), and that GSDMD killed WT and Δ*clsA S. flexneri* with similar efficiency (Figures 6A-6C and S6B-S6E). These data show that bacterial cardiolipin is not required for GSDMD-dependent macrophage killing of *S. flexneri*. Emerging evidence indicates that GSDMD can form pores in cardiolipin-enriched mitochondrial membranes, which triggers mitochondrial dysfunction and leads to release of reactive oxygen species (ROS).^48,55–57^ Although mitochondrial ROS have been proposed to kill intracellular bacteria,^3,58,59^ they are not responsible for macrophage killing of *S. flexneri*, as bacterial killing was not impacted by ROS inhibition (Figure S5A). Together, these findings uncover a previously undescribed cardiolipin-independent mechanism of GSDMD targeting and killing of intracellular bacteria.

Ectopic expression of the N-terminal pore-forming domain of GSDMD (NT-GSDMD) was sufficient to kill intracellular *S. flexneri* (Figure 5E). As NT-GSDMD has been shown to kill the gram-positive bacteria *Staphylococcus aureus* and *Listeria monocytogenes*, and protoplasts of *Escherichia coli* and *Bacillus megaterium*,^17,19^ all of which lack an outer membrane, we speculate that GSDMD killing of *S. flexneri* requires pore formation in the bacterial inner membrane. How GSDMD gains access to the inner membrane remains unknown. GSDMD may recruit auxiliary host factors that destabilize the bacterial membrane. A candidate molecule would be GBP1, which destabilizes the bacterial membrane, and under certain conditions, contributes to bacterial lysis;^43^ however, we found that GBP1 is dispensable for killing *S. flexneri* (Figure 4E), indicating that it is not required for GSDMD-dependent bacterial restriction. As NT-GSDMD is efficient at killing *S. flexneri* Δ*clsA* (Figures 6D-6E and S6G), we hypothesize that GSDMD targets alternative bacterial lipids or surface determinants.

In human macrophages, *S. flexneri* activates both NLRP3 and NLRC4 inflammasomes.^8,10^ We found that NLRC4, but not NLRP3, is essential for intracellular bacterial killing (Figures 3E-3G and 7A). Given that NLRP3 responds to disturbances in cellular homeostasis,^60–62^ we speculate that NLRP3 activation instead functions as a secondary pathway that promotes pyroptosis and downstream bacterial clearance by other mechanisms, including non-self-autonomous mechanisms, when macrophage-intrinsic mechanisms fail to control the infection.

In human macrophages, *S. flexneri* also activates caspase-4.^9^ Whereas caspase-4 contributes to control of *Salmonella* primarily at late stages of infection,^22^ we found that caspase-4 restricts *S. flexneri* early during infection, although only in IFNγ-primed macrophages (Figure 4B). Importantly, upon IFNγ priming, both caspase-1 and caspase-4 contribute to restriction of *S. flexneri* survival (Figure 4C), demonstrating dual contribution of canonical and non-canonical inflammasome pathways to controlling intracellular bacteria. IFNγ priming was associated with increases in levels of both caspase-1 and caspase-4, which was associated with increased restriction of intracellular *S. flexneri* (Figures 4C, 4G, and 7B) and reduced dependence on upstream pattern-recognition receptors (Figures 3H, and 4E-4F). The increased levels of these caspases are presumably due to upregulation of expression of *CASP1* and *CASP4*.^29,30^ Increased levels of the caspase in turn lead to enhanced activation of downstream GSDMD (Figure 4G), resulting in increased killing of bacteria (Figure 7B).

**Figure 7.**
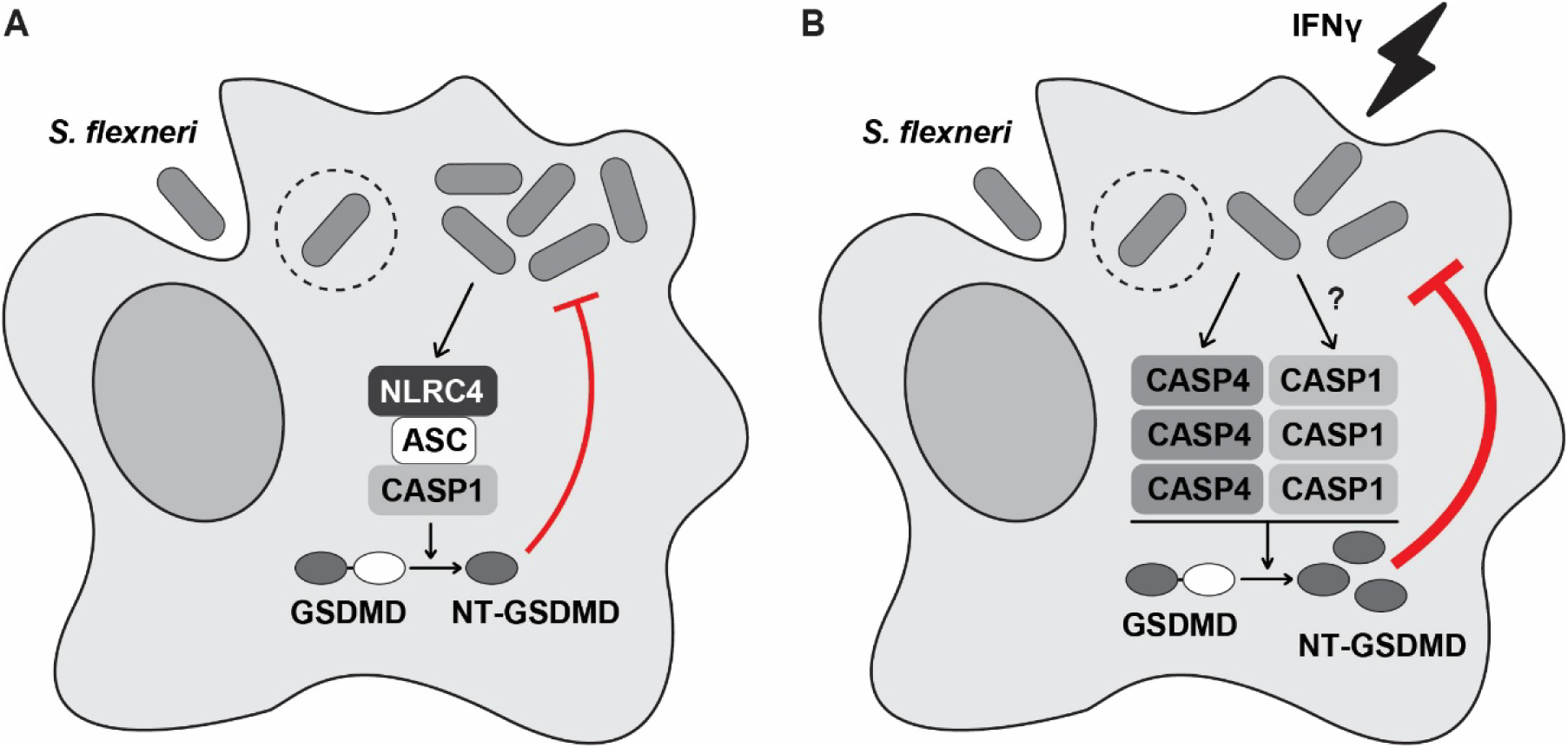
Model of inflammasome-based restriction of *S. flexneri* in human macrophages. (A) First contact of macrophages with pathogens occurs in the absence of IFNγ priming. In this setting, *S. flexneri* is recognized by NLRC4, which leads to processing and activation of caspase-1 (CASP1) and downstream GSDMD. GSDMD targets and kills intracellular *S. flexneri*. (B) Once the innate immune system is activated, macrophages are primed with IFNγ,^53^ leading to increased caspase-4 (CASP4) and caspase-1 (CASP1) and enhanced activation of GSDMD, which increases killing of intracellular *S. flexneri*. In both cases, GSDMD kills *S. flexneri* independently of bacterial cardiolipin and host cell lysis.

IFNγ polarizes macrophages towards the classically activated, pro-inflammatory M1 state.^27,28^ Here, we identify a previously unrecognized pathway of inflammasome-dependent control that contributes to the pro-inflammatory program of M1 macrophages. In addition, IFNγ induces several complementary bactericidal pathways, including the RIG-I pathway that has been associated with restriction of *S. flexneri* replication,^37^ and apolipoprotein L3, which directly targets bacteria for lysis.^36^ Although discovered in mouse embryonic fibroblasts and HeLa cells, we do not exclude potential contributions of these pathways to restriction of *S. flexneri* in human macrophages; either or both could be responsible for the observed IFNγ-dependent GSDMD-independent restriction of *S. flexneri* (Figures 3A and S3C).

*S. flexneri*-infected macrophages produce IL-12 that induces production of IFNγ by NK cells, Nkp46^+^ innate lymphoid cells (ILCs), and γδT cells,^26^ which has been proposed to enhance inflammasome activation in epithelial cells of the infected intestinal mucosa and thereby promote host defense. In addition, we found previously that IFNγ and NK cells are critical for clearance of *S. flexneri* in mouse models of infection, including in mouse macrophages.^63^ Here, we show that in human macrophages, IFNγ contributes to increased bacterial clearance via a macrophage-intrinsic inflammasome-dependent mechanism.

In conclusion, our work demonstrates a previously undescribed mechanism of GSDMD-mediated killing of cytosolic bacteria that is independent of bacterial cardiolipin, identifying a previously unrecognized mechanism of microbial recognition. GSDMD is required for macrophage killing of *S. flexneri* in IFNγ-independent and IFNγ-dependent pathways, with both the canonical and the non-canonical inflammasomes contributing in the presence of IFNγ. As such, our work describes a new cell-autonomous defense program of human macrophages against cytosolic bacteria and broadly contributes to our understanding of innate immunity against intracellular pathogens.

## MATERIALS AND METHODS

### Bacterial strains and growth conditions

All *Shigella* strains used in this study are isogenic derivatives of *S. flexneri* wild-type serotype 2a strain 2457T.^64^ The Δ*ipaC* and Δ*ipaH9.8* mutants have been described.^65,66^ The Δ*clsA* mutant (cardiolipin-deficient) was a gift from S. Payne.^52^ All bacteria were grown in tryptic soy broth at 37°C with aeration upon seeding from Congo red agar plates.

### Mammalian cell culture

Human-derived THP-1 monocytes (TIB-202) were obtained from American Type Culture Collection (ATCC) and cultured in RPMI 1640 medium (Gibco, 11875119) supplemented with 10% (v/v) heat-inactivated fetal bovine serum (FBS, heat-inactivated, sterile-filtered, R&D systems) and 10 mM HEPES (Gibco, 15630-080). HEK293T cells (CRL-3216) were obtained from ATCC and cultured in Dulbecco’s modified Eagle’s medium (DMEM, Gibco, 11965-092) supplemented with 10% (v/v) heat-inactivated FBS (R&D systems). All cells were grown in 5% CO_2_ in a humidified incubator at 37°C.

### Cell treatments

To induce differentiation into macrophages, THP-1 monocytes were incubated with 100 ng/mL phorbol 12-myristate 13-acetate (PMA, Sigma-Aldrich, P8139) for 24 hours, after which they were rested in media without PMA overnight prior to use in experiments. Where indicated, macrophages were primed with 10 ng/mL gamma interferon (IFNγ, human, PeproTech 300-02-100UG) for 16 to 20 hours. Where indicated, macrophages were treated with glycine at 5 mM and disulfiram (Tocris, 3807) at 30 µM, respectively, for 1 hour prior to infection. For inhibition of NLRP3 activation, macrophages were pre-treated with MCC950 (InvivoGen, inh-mcc) at indicated concentrations for 1 hour prior to infection or nigericin treatment, and treated continuously after. For activation of the NLRP3 inflammasome, macrophages were primed with 0.5 µg/mL LPS (0111:B4 from *E. coli*, InvivoGen, tlrl-3pelps) for 3 hours and then treated with 10 µM nigericin (InvivoGen, tlrl-nig) for 90 mins. To activate NLRP3 in *S. flexneri*-infected cells, macrophages were primed with LPS for 2 hours prior to infection; at 1 hour 15 mins of infection, macrophages were treated with 20 µM nigericin for 2 to 2.5 hours. Pam3CSK4 (InvivoGen, tlrl-pms) was used at 1 µg/mL overnight to prime inflammasome activation. To inhibit oxidative stress, macrophages were treated with reduced L-glutathione (Sigma-Aldrich, G6013) at 10 mM for 1 hour prior to infection.

### Bacterial infections

Single red colonies of *S. flexneri* were used to inoculate overnight cultures. PMA-differentiated macrophages or HEK293T cells were washed with serum-free media and infected at an MOI of 10 with *S. flexneri* grown to exponential phase after back-dilution from the overnight culture. Bacterial inoculum was prepared in media without serum. Bacteria were centrifuged onto cells at 800 g for 10 min at room temperature, followed by incubation at 37°C. After 30 min, cells were washed twice with Hanks’ Balanced Salt Solution (HBSS, Gibco, 14025-092) and serum-free media containing gentamicin at 25 µg/mL was added to kill extracellular bacteria. Samples for LDH release assay and western immunoblot were collected at 3 hours of infection.

### Quantification of bacteria

To quantify bacteria in attached cells, cells were infected, as above, in 48-well dishes containing 2x10^5^ cells per well. Media with gentamicin was removed after 30 mins, cells were washed twice with HBSS and overlayed with fresh serum-free media. At 3 hours of infection, cells were washed three times with HBSS and lysed with 0.5% Triton X-100 for 5 mins at room temperature. The lysates were serially diluted and plated on agar plates for enumeration of bacteria. To quantify bacteria in the “media” fraction containing detached cells and released bacteria, cell culture supernatants and HBSS washes collected in the same dishes at 3 hours of infection were centrifuged at 8000 g for 2 mins. The pellets were lysed with 0.5% Triton X-100 and bacteria were enumerated as above. To determine efficiency of bacterial invasion, cells were washed three to five times with HBSS following invasion and lysed as above.

### Vacuolar escape

To assess the ability of *S. flexneri* to escape the pathogen-containing vacuoles, macrophages were infected as above. Following invasion, macrophages were washed three times with HBSS and overlayed with media containing 25 µg/mL gentamicin, or 25 µg/mL gentamicin and 200 µg/mL chloroquine (Sigma-Aldrich, C6628). After 1 hour, macrophages were lysed and bacteria were enumerated as above.

### Macrophage death

Macrophage cell death was assessed by release of LDH due to loss of plasma membrane integrity. 50 μL of cell culture supernatants from a 96-well plate were collected at 3 hours of infection and CyQUANT™ LDH Cytotoxicity Assay (ThermoFisher Scientific, C20301) was used following the manufacturer’s instructions.

### Protein extraction and western immunoblot analysis

To detect released caspases, 100 μL of cell culture supernatants from a 48-well plate were collected at 3 hours of infection or 90 minutes of nigericin treatment and mixed with 35 μL of 4x Laemmli buffer [250 mM Tris (pH 6.8), 8% SDS, 50% glycerol, 0,04% bromophenol blue] supplemented with cOmplete EDTA-free protease inhibitor (Roche, 11873580001). To detect cell-associated proteins, cells in the same plate were lysed directly in 100 μL of 4x Laemmli buffer supplemented with the same protease inhibitor. Following SDS-polyacrylamide gel electrophoresis (SDS-PAGE), proteins were transferred onto a nitrocellulose membrane.

Antibodies used for immunoblotting were as follows: caspase-1 (Abcam, ab207802) rabbit monoclonal antibody at 0.5 μg/mL (1:1000), GBP1 (Abcam, ab131255) rabbit monoclonal antibody at 0.25 μg/mL (1:5000), caspase-4 (Santa Cruz, sc-56056) and GSDMD (Santa Cruz, sc-81868) mouse monoclonal antibodies at 0.5 μg/mL (1:200). Secondary antibodies were goat anti-rabbit immunoglobulin G (IgG) or goat anti-mouse IgG, each conjugated to horseradish peroxidase (HRP) (Jackson ImmunoResearch). Antibodies specific for β-actin were HRP-conjugated (Sigma, A3854). Immunoreactive bands were visualized by chemiluminescence with SuperSignal West Pico PLUS or SuperSignal West Femto Maximum Sensitivity substrates (Thermo Fisher Scientific, 34580 and 34096).

### Generation of CRISPR deletions

Chemically modified sgRNAs (Synthego) were electroporated as ribonucleoprotein (RNP) complexes into freshly cultured THP-1 monocytes using the Neon Transfection System (Invitrogen, MPK5000) following the manufacturer’s instructions. 100 pmol of specific sgRNAs (Table S1) were mixed with 25 pmol (4:1 ratio) or 11 pmol (9:1 ratio) of SpCas 2NLS nuclease (Synthego) in Resuspension Buffer R (Invitrogen, MPK1069B) and incubated for 15 minutes at room temperature. THP-1 monocytes were washed in PBS and resuspended in Buffer R. The RNP complexes were electroporated by pulsing twice 1x10^6^ cells in the 10 µL Neon Tip at 1700 V and 2 ms pulse width. Electroporated cells were resuspended in 2.5 mL complete growth medium in a 6-well plate and incubated at 37°C. After 72 hours, 1.5 mL cell suspension was transferred to a 75 cm^2^ culture flask, and 1 mL was used to purify genomic DNA using DNeasy Blood and Tissue kit (Qiagen). Following PCR and sequencing, editing efficiency was determined using ICE Analyzer (Synthego). The cells in culture flask were limiting-diluted and expanded as single clones in a 96-well dish. Isolated single clones were validated for homozygous out-of-frame deletions by sequencing.

### RNA interference

PMA-differentiated THP-1 macrophages were transfected with siRNAs. THP-1 monocytes were seeded in a 48-well dish at 2x10^5^ cells per well and differentiated into macrophages with 100 ng/mL PMA. After 24 hours, the media was replaced with fresh media without PMA. The cells were transfected with GBP1-specific Silencer^TM^ Select Pre-Designed siRNAs (Invitrogen, 4392420, siRNA ID: s5620 and s5621) at 50 nM using Lipofectamine 3000 (Invitrogen, L3000008) following the manufacturer’s instructions. For scrambled siRNA (negative control), AllStars Negative Control siRNAs (Qiagen) were used. After 24 hours, the cells were primed with 10 ng/mL IFNγ without replacing the media. After 16 to 20 hours, samples were collected for western immunoblot, or cells were infected for quantification of bacteria.

### GSDMD killing assay

The pcDNA3.1 vector expressing full-length GSDMD (FL GSDMD) and pLV-3xFLAG vector expressing the N-terminal pore-forming domain of GSDMD (NT-GSDMD) were a gift from H. Wu. The coding sequence of NT-GSDMD was subcloned into pcDNA3.1 using primers 5’ AGATCTGCTAGCATGGACTACAAAGACC (forward) and 5’ AGATCTCTCGAGCTAATCTGTCAGGAAGTTG (reverse) and NheI and XhoI restriction enzymes (New England Biolabs). The sequence of the generated pcDNA3.1 expressing NT-GSDMD was confirmed by whole plasmid sequencing.

HEK293T cells were seeded in a 48-well dish at 5x10^4^ cells per well. After 24 hours, cells were transfected with 150 ng of an empty pcDNA3.1 vector or pcDNA3.1 vector expressing FL GSDMD or NT-GSDMD, using FuGENE® 6 (Promega, E269A) following the manufacturer’s instructions. After 18 hours, samples were collected for western immunoblot, or cells were infected for quantification of bacteria in attached cells and the “media” fraction, as above.

### GSDMD subcellular localization

THP-1 monocytes were seeded at 4x10^5^ cells per well onto microscope coverslips in a 24-well dish. The cells were differentiated into macrophages, primed with IFNγ, and infected with *S. flexneri* as above. At 50 mins of infection, cells were washed three times with HBSS, fixed with 2% paraformaldehyde for 15 mins, washed with PBS, permeabilized with 0.2% Triton X-100 (in PBS) for 10 mins, and washed with PBS. To limit nonspecific antibody binding, cells were blocked with 5% bovine serum albumin (BSA) (in PBS) overnight at 4°C. Cells were incubated with a rabbit polyclonal antibody to GSDMD (Proteintech, 20770-1-AP), used at 3.2 μg/mL (1:250), for 2 hours, and washed with PBS. All subsequent steps were performed in the dark. Cells were incubated with a goat anti-rabbit IgG conjugated to Alexa Fluor 594 (Thermo Fisher, A11012), used at 2 μg/mL (1:500), for 50 mins, washed extensively with PBS, and incubated with a rabbit polyclonal antibody to *Shigella* (Virostat, 0903), used at 16 μg/mL (1:300), for 45 mins. Cells were incubated with Hoechst 33342 (Invitrogen, H3570) at 1 μg/mL for 15 mins. After washing with PBS, coverslips were mounted onto microscope slides using Prolong^TM^ Diamond Antifade Mountant (Invitrogen, P3670), and the samples were left to dry overnight. Cells were imaged using a Nikon Eclipse TE300 inverted fluorescence phase contrast microscope.

### Statistical analysis

Unless otherwise indicated, all reported data are from at least three independent biological replicates. For bar graphs, each biological replicate is indicated as a separate symbol. GraphPad Prism10 software was used for data graphing and statistical analyses. A normal-based 95% confidence interval (mean ± 2 SD) was applied for statistical assessment. Details of the statistical tests used are provided in the figure legends.

## ACKNOWLEDGMENTS

We thank Shelley Payne for providing the cardiolipin-deficient (Δ*clsA*) mutant of *S. flexneri*, Hao Wu for providing the pcDNA3.1 vector expressing full-length human GSDMD and the pLV-3xFLAG vector expressing NT-GSDMD. This work was supported by NIH T32 AI007061 (to M.S.), F32 AI188955 (to M.S.), and R01 AI173030 (to M.B.G.).

