## Supplemental information for "Dual NLRC4 and non-canonical inflammasome signaling drives human GSDMD-mediated killing of *Shigella flexneri* independently of bacterial cardiolipin"

**SUPPLEMENTAL DATA**

**Table S1. Sequences of sgRNAs used in this study.**

|  | **Sequence** | **Gene target** | **sgRNA:Cas9 ratio** |
| --- | --- | --- | --- |
| sgRNA 1 | 5’ UUCCCCUCCUCAGGAGCAUG | *GSDMD* | 9:1 |
| sgRNA 2 | 5’ GACCCCAUGCUCCUGAGGAG |  |  |
| sgRNA 3 | 5’ CACCGACCAGCCUGCAGAGCUCCAC |  |  |
| sgRNA 4 | 5’ AAACGUGGAGCUCUGCAGGCUGGUC |  |  |
| sgRNA 5 | 5’ CUAAACAGACAAGGUCCUGA | *CASP1* | 4:1 |
| sgRNA 6 | 5’ AAGCUGUUUAUCCGUUCCAU |  |  |
| sgRNA 7 | 5’ AAACAUCAUUUGCUGCGAGA | *NLRC4* | 4:1 |
| sgRNA 9 | 5’ CAUUCCCAUUCUUUGAAUAA |  |  |
| sgRNA 10 | 5’ CUCCUCAGUGAAUUUCAUAA |  |  |
| sgRNA 11 | 5’ GAGAAGCAAGAUGGCAGAAU | *NLRP11* | 4:1 |
| sgRNA 12 | 5’ CGUGUUGCCAAUCUCUUAUG |  |  |
| sgRNA 13 | 5’ GUGUUGCCAAUCUCUUAUGA |  |  |
| sgRNA 14 | 5’ UGCGUAAGGAAGAUCUUUGUAGG |  |  |

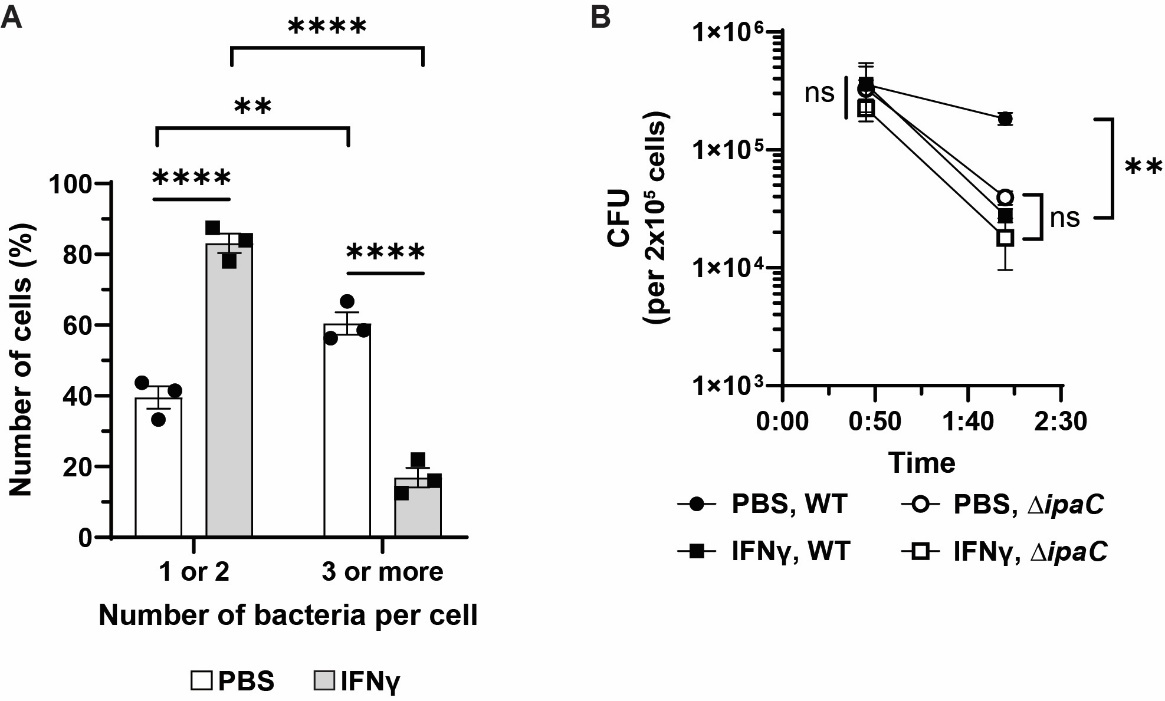

**Figure S1. Supplemental data to Figure 1 (main text).**

(A) Supplemental information to Figure 1B (main text). IFNγ induced reduction in the number of bacteria per macrophage. Minimum of 360 cells (120 per biological replicate) were scored for each condition. Macrophages were infected for 2 hours.

(B) Supplemental information to Figure 1F (main text). Reduction in intracellular *S. flexneri* over time in macrophages with or without IFNγ priming.

Data represent the mean ± SEM. ***p* < 0.01, *****p* < 0.0001, ns, not significant, by ordinary two-way ANOVA (A) or two-tailed unpaired Student’s t-test (B).

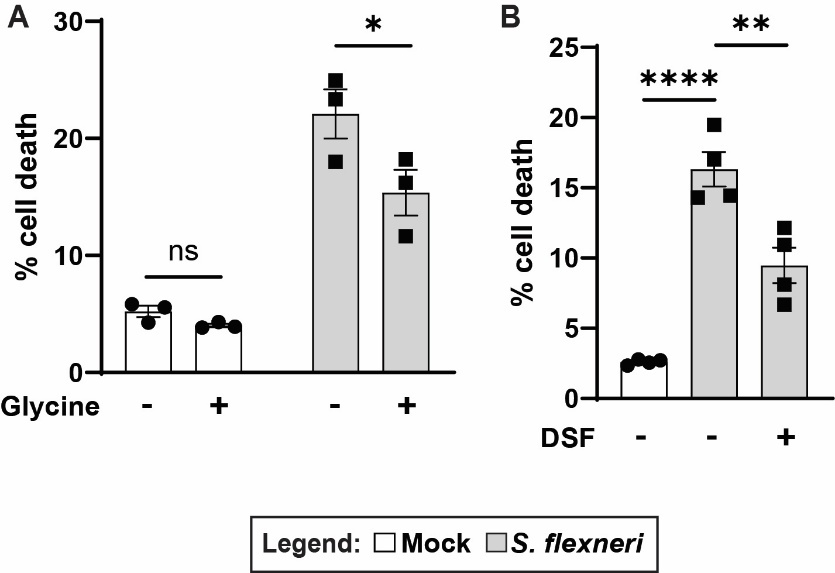

**Figure S2. Impact of inhibition of GSDMD pore formation and plasma membrane rupture on macrophage death.**

(A-B) Cell death (measured as LDH release) of macrophages infected with *S. flexneri* for 3 hours, with or without addition of glycine (an inhibitor of plasma membrane rupture) (A), or with or without addition of disulfiram (DSF, an inhibitor of GSDMD pore formation) (B).

Data represent the mean ± SEM. **p* < 0.05, ***p* < 0.01, *****p* < 0.0001, ns, not significant, by ordinary one- (B) or two-way (A) ANOVA.

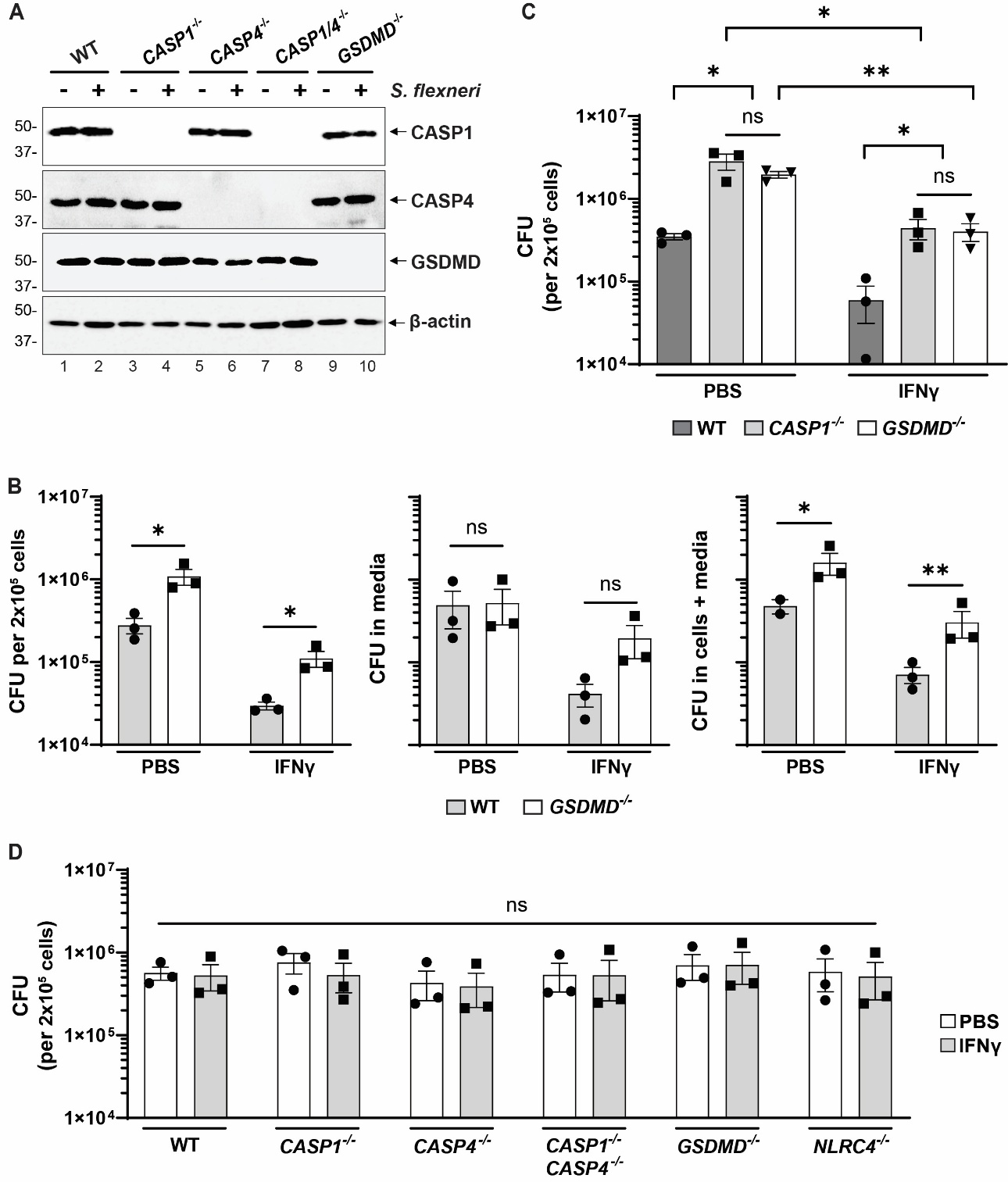

**Figure S3. Supplemental data to Figure 3 (main text).**

(A) Absence of inflammasome proteins in indicated knockout macrophages. Representative immunoblots.

(B) Rescue of intracellular *S. flexneri* in absence of GSDMD is not due to GSDMD-dependent bacterial loss in media or detached macrophages. Graphed are *S. flexneri* quantified in attached macrophages (left graph), cell culture supernatants and washes (media), which contain detached macrophages and released bacteria (middle graph), and both combined (right graph).

(C) Rescue of intracellular *S. flexneri* in the absence of *CASP1* or *GSDMD*, with or without IFNγ priming.

(D) Lack of impact of knockout of the indicated inflammasome proteins on the efficiency of *S. flexneri* invasion into macrophages, assayed at 45 min. of infection.

Data represent the mean ± SEM. **p* < 0.05, ***p* < 0.01, ns, not significant, by two-tailed unpaired Student’s t-test (B) or ordinary two-way ANOVA (C-D).

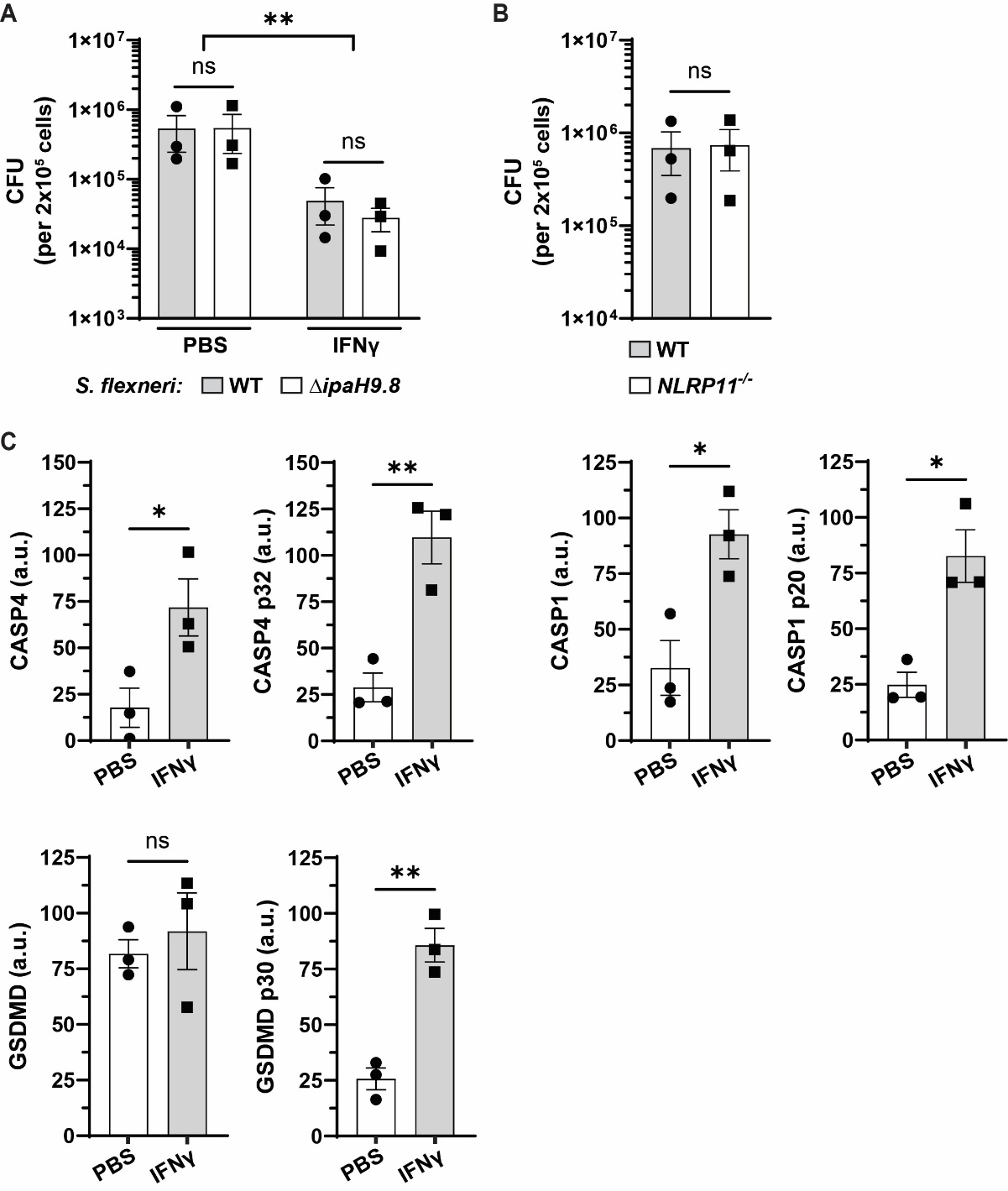

**Figure S4. Supplemental data to Figure 4 (main text)**.

(A) *S. flexneri* Δ*ipaH9.8* is restricted by macrophages similar to WT *S. flexneri*. Macrophages were infected for 3 hours.

(B) Deletion of macrophage *NLRP11* does not impact efficiency of *S. flexneri* invasion, determined at 45 min. of infection.

(C) Quantification (band densitometry) of western immunoblots from Figure 4G (main text), measuring release of cleaved caspase-4 (CASP4, top left graphs) and caspase-1 (CASP1, top right graphs) into cell culture supernatants, and cleaved GSDMD in cell lysates (bottom graphs) in macrophages infected for 3 hours. Protein levels, expressed in arbitrary units (a.u.), were normalized to β-actin.

Data represent the mean ± SEM. **p* < 0.05, ***p* < 0.01, ns, not significant, by ordinary two-way ANOVA (A) or two-tailed unpaired Student’s t-test with Welch’s correction (B and C).

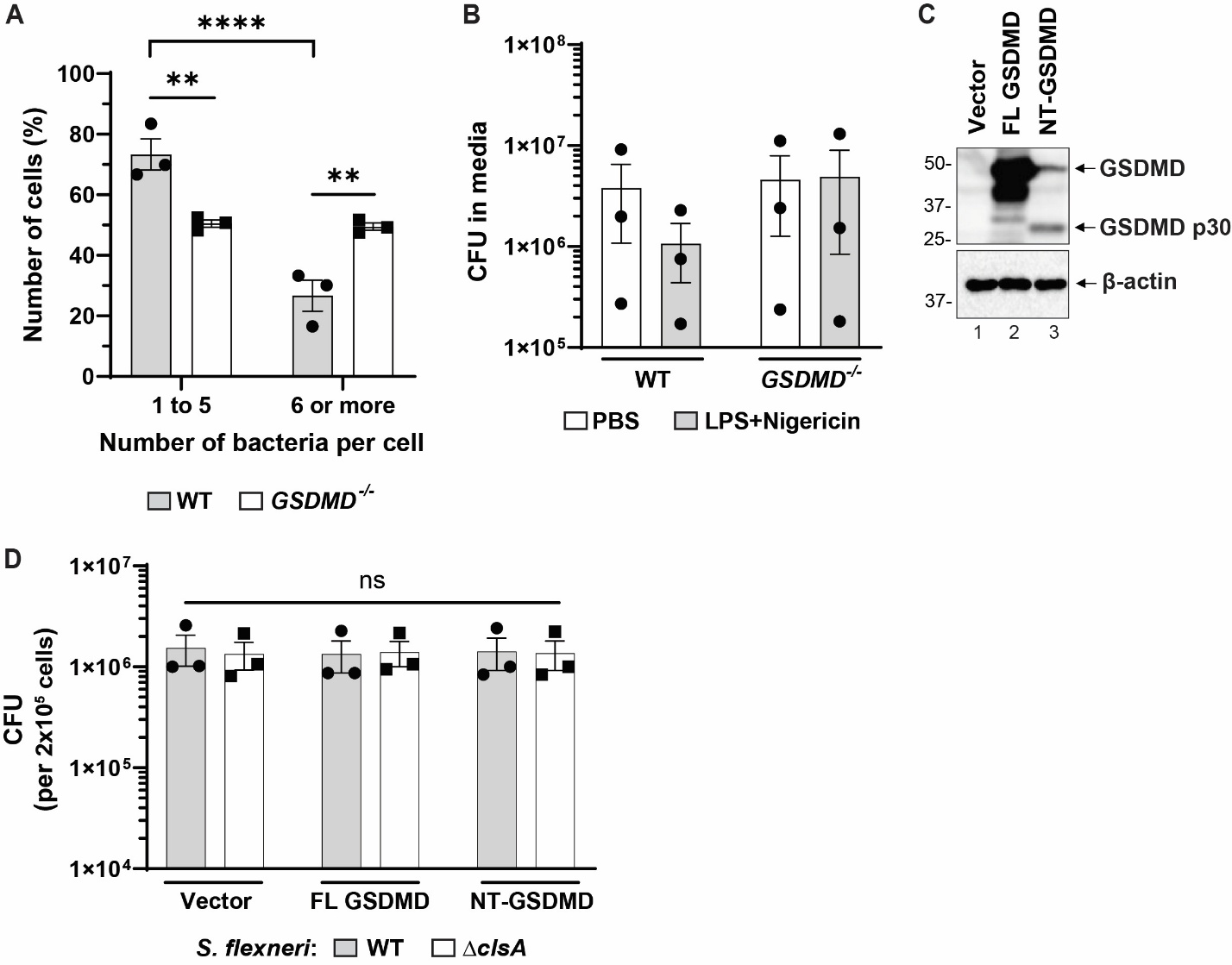

**Figure S5. Supplemental data to Figure 5 (main text).**

(A) Supplemental information to Figure 5B (main text). GSDMD dependence of the number of bacteria per macrophage. Minimum of 480 cells (160 per biological replicate) were scored for each condition. Macrophages were infected for 2 hours.

(B) Enhanced killing of *S. flexneri* by treatment with extracellular LPS and nigericin is not due to bacterial loss into cell culture supernatants. *S. flexneri* quantified in cell culture supernatants and washes (media), which contain detached macrophages and released bacteria.

(C) Representative immunoblot showing expression of indicated GSDMD constructs in HEK293T cells.

(D) Lack of difference in efficiency of invasion by WT and Δ*clsA S. flexneri* into HEK293T cells expressing indicated GSDMD constructs. Assayed at 45 min. of infection.

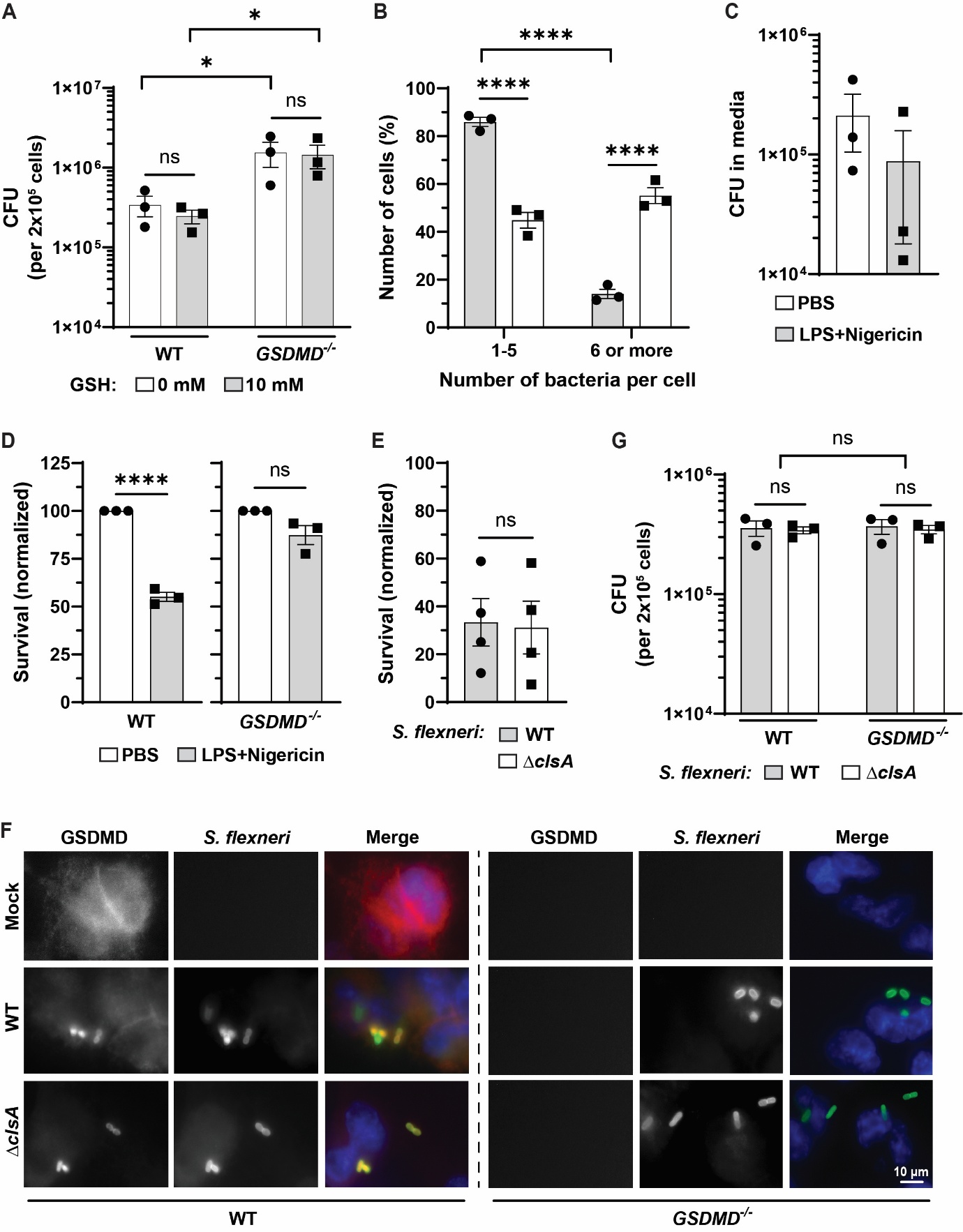
**Figure S6. GSDMD kills *S. flexneri* independently of oxidative stress and bacterial cardiolipin (supplemental data).**

(A) Impact of oxidative stress, with or without IFNγ priming, on GSDMD-mediated killing of *S. flexneri*. Bacterial counts following treatment of indicated macrophages with the oxidative stress inhibitor reduced glutathione (GSH).

(B) Supplemental information to Figure 6B (main text). IFNγ priming-dependent reduction in the number of *S. flexneri* Δ*clsA* per macrophage. Minimum of 420 cells (140 per biological replicate) were scored for each condition. Macrophages were infected for 2 hours.

(C) Enhanced killing of *S. flexneri* Δ*clsA* by treatment with extracellular LPS and nigericin is not due to bacterial loss into cell culture supernatants. Bacteria quantified in cell culture supernatants and washes (media), which contain detached macrophages and released bacteria.

(D) GSDMD is required for enhanced killing of *S. flexneri* Δ*clsA* upon treatment with extracellular LPS and nigericin. For each bacterial strain, survival is normalized to macrophages not treated with extracellular LPS and nigericin.

(E) WT and Δ*clsA* *S. flexneri* survive similarly following treatment with extracellular LPS and nigericin. For each bacterial strain, survival is normalized to macrophages not treated with extracellular LPS and nigericin.

(F) Colocalization of GSDMD and *S. flexneri*. Immunofluorescence at 50 mins. of infection with WT or Δ*clsA* *S. flexneri*, with or without IFNγ priming, using antibodies to GSDMD (red) and *S. flexneri* (green).

(G) Lack of difference in efficiency of invasion by WT and Δ*clsA S. flexneri* into macrophages. Assayed at 45 min. of infection.

Data represent the mean ± SEM. **p* < 0.05, *****p* < 0.0001, ns, not significant, by two-tailed unpaired Student’s t-test (C-E) or ordinary two-way ANOVA (A-B, G).
